# Chromosome disentanglement driven via optimal compaction of loop-extruded brush structures

**DOI:** 10.1101/616102

**Authors:** Sumitabha Brahmachari, John F. Marko

## Abstract

Eukaryote cell division features a chromosome compaction-decompaction cycle that is synchronized with their physical and topological segregation. It has been proposed that lengthwise compaction of chromatin into mitotic chromosomes via loop extrusion underlies the compaction-segregation/resolution process. We analyze this disentanglement scheme via considering the chromosome to be a succession of DNA/chromatin loops - a polymer “brush” - where active extrusion of loops controls the brush structure. Given topoisomerase (TopoII)-catalyzed topology fluctuations, we find that inter-chromosome entanglements are minimized for a certain “optimal” loop that scales with the chromosome size. The optimal loop organization is in accord with experimental data across species, suggesting an important structural role of genomic loops in maintaining a less entangled genome. Application of the model to the interphase genome indicates that active loop extrusion can maintain a level of chromosome compaction with suppressed entanglements; the transition to the metaphase state requires higher lengthwise compaction, and drives complete topological segregation. Optimized genomic loops may provide a means for evolutionary propagation of gene-expression patterns while simultaneously maintaining a disentangled genome. We also find that compact metaphase chromosomes have a densely packed core along their cylindrical axes that explains their observed mechanical stiffness. Our model connects chromosome structural reorganization to topological resolution through the cell cycle, and highlights a mechanism of directing Topo-II mediated strand passage via loop extrusion driven lengthwise compaction.

---

Chromosomes are biopolymer structures made up of long, multi-megabase-pair DNAs bound by proteins, residing in confined spaces. For eukaryotes, the nucleus is the confining compartment, while bacterial chromosomes are confined by the cell itself. Without their organization into looped structures, the confinement would lead to cellular chromosomes forming an entangled, semi-dilute polymer solution[1], i.e., the fraction of the total confinement volume occupied by all the genomic segments (average volume fraction) is high enough to force strong overlap between different chromosomes (simply considered as linear polymers). However, as we argue in the following, the organization of chromosomes into loops with a “polymer-brush”-like architecture [Fig. 1(b)] leads to lower inter-chromosomal overlap for a fixed volume fraction.

**FIG. 1.**
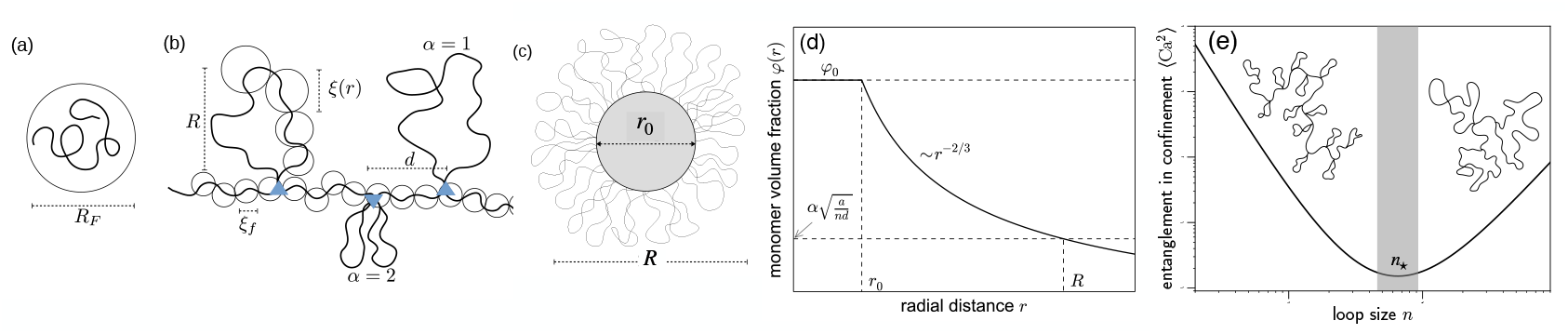
(a) Flory radius or equilibrium dimension of a self-avoiding polymer [Eq. A1]. (b) Sketch of loop-extruded chromosome showing chromatin loops connected by a backbone, where circles represent blobs of size *ξ*, that depends on the radial distance from the backbone. Out of the three schematic loops sketched, the middle one is divalent (*α* = 2) and the other two are monovalent (*α* = 1). (c) Cross-sectional view of compacted chromosome. Circular cross-section corresponding to the gray-shaded area is the densely-packed-chromatin core of width *r*_0_. (d) Monomer volume fraction *φ*(*r*) is maximum inside the core [*φ*_0_ ≈ 1] and decays radially outwards. (e) For a fixed genome size *N*, backbone size *m*, and nuclear volume, inter-chromosome entanglements per chromosome 〈Ca^2^〉 [(4)] shows a minimum for an optimal configuration with loop size *n*_*_ (shown by a shaded region). This follows from the geometry and finite size of chromosomes. Schematic pictures of finite-size chromosomes in the small- and large-loop regimes are shown.

An important question in chromosome biopphysics is how can topology-manipulating enzymes: Type-II DNA topoisomerases (Topo II), that catalyze the passage of one genomic segment through another, drive global disentanglement of chromosomes. Since individual Topo IIs cannot sense the global entanglement topology of chromosomes, we consider Topo II to facilitate random strand passage. This lets the chromosomes pass through each other akin to a “phantom” polymer chain where the inter-chromosomal topology fluctuates [2]. However, random strand passage is not enough to completely disentangle long linear polymers, because the entropically favored state is the one with higher inter-chromosomal mixing or volume overlap. Modeling chromosomes as arrays of polymer loops, we find that the compaction generated from chromosomal loop organization is capable of driving inter-chromosome disentanglement and segregation, under the conditions of fluctuating topology. Our results are complementary to, and to some extent establish a theoretical description of recent simulation results that show how loop-extruding protein machines are able to geometrically and topologically organize long polymer-like chromosomes [3–5].

Eukaryote chromosomes undergo significant and highly ordered compaction during mitosis. This process cannot be “condensation” in the usual sense of that term: uncontrolled self-adhesion of chromatin will lead to a compact and highly entangled genomic globule. More plausibly, chromosome segregation is based on “lengthwise compaction” simultaneous with Topo II-mediated topology changes, cooperating to drive progressive physical and topological segregation [6, 7]. Processive extrusion of genomic loops by loop-extruding enzymes has been proposed to underlie lengthwise compaction [3, 8] and the formation of the long-observed cylindrical brush (loop array) structure of metaphase chromosomes [9–11]. The “loop-extrusion” hypothesis proposes a microscopic mechanism to achieve length-wise compaction of chromosomes based on molecular-motor-generated tension along the polymer contour [3, 8]. In other models of chromosome compaction, such a compaction-generating tension may be effectively generated by mechanisms like supercoiling flux arising from transcription and replication [12], or “sliding” of genomic contacts driven by directed motion of molecular slip-links [13, 14], or diffusion of genomic segments in a hypothesized data-driven potential [15, 16].

Structural Maintenance of Chromosomes (SMC) complexes are thought to be responsible for organizing chromosome structure [17–25], and recently have been directly observed to processively translocate and loop-extrude DNA[26, 27]. The three-dimensional conformation of the interphase genome has been observed to be organized into loops that are associated with regulation of gene expression, and these loops appear to be actively driven by a concerted action of proteins like SMC complexes and other architectural proteins [20, 24, 25, 28–33]. Electron microscope images of metaphase chromosomes have directly observed the loop organization, supporting a polymer brush model with radially emanating loops [9, 10, 34].

We model chromosomal DNA/chromatin as a long polymer, and chromosomes as a polymer brush-like steady state structure where the “bristles” represent chromatin loops. Comparing with experimentally observed genomic loops, we find that chromosomes appear to be organized into “optimal”-sized loops, that maximizes compaction, and simultaneously minimizes inter-chromosome entanglement. Qualitatively, the optimal size occurs since for small loops, chromosomes have a long axial length and easily become highly entangled with other chromosomes. As the loops grow in size, the chromosome is gradually lengthwise-compacted to become a cylindrical brush of size smaller than the original chromosome [Eq. 2], and entanglements between chromosomes are reduced. If the loops become so large that they are comparable to the size of the chromosomes themselves, the chromosome becomes a “star polymer” with long bristles which once again become highly entangled with bristles of neighbor chromosomes [Fig. 1(d)]. While we will discuss our results primarily in the context of eukaryotes, we also show how they can be applied to bacterial chromosomes.

Here, we show that loop-extruding motors, capable of exerting forces in the picoNewton (pN) range, are sufficient to drive compaction to a stage where osmotic repulsion between chromosomes will lead to a disentangled chromosomes, given that entanglement topology is allowed to change (Topo II is active). Compaction tension generates stress in chromosomes, which is then relaxed through entanglement release by Topo II: the coupling of compaction to topological simplification provides the key driving force directing Topo II to disentangle the genome.

## I. POLYMER MODEL AND METHODS

We consider chromatin (or DNA coated with proteins) as a long self-avoiding polymer that is organized as an array to loops that resembles a “bottle brush” polymer. The brush is in a constrained local thermal equilibrium, because maintaining the brush structure requires continual work to be done by SMC complexes that secure the loop anchors against thermal agitation. The motor activity of SMC complexes leads to active reorganization of the chromosome polymer, but since the self-organization time is much longer than the equilibration (Rouse) time of a typical loop even including entanglement release dynamics by Topo II (see Discussion) we may employ static polymer scaling laws.

We implement a two-level polymer physics model: at shorter length scales there is chromatin behaving as a self-avoiding polymer, then, at longer lengths –comparable to chromosome radii, where inter-chromosomal interactions dominate– we consider chromosomes as effective polymers made up of cylindrical brush segments. Our chromatin is a chain of spherical monomers of diameter *a*, such that a chromatin chain of *N* monomers “swells” up due to self-avoiding monomer correlations, and assumes a dimension given by its “Flory radius”:

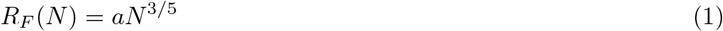

We make the choice of monomer as one nucleosome (200 bp DNA): *a* = 10 nm. Recent electron-microscopy [35–37], and super-resolution imaging analyses [38] are in line with a highly flexible chromatin fiber model with each nucleosome as a polymer segment, rather than a more traditional “stiff 30 nm fiber” model. In the bacterial case, the polymer is DNA coated by nucleoid-associated proteins with a similar value of *a* but containing 150 bp DNA. The cylindrical aspect ratio of the bacterial “chromatin” monomer is not critical to our results.

When multiple chromatin chains (say, *k* chains, such that the genome size is 𝒢 = *Nk*) are confined within a volume *V* « *kR*_*F*_ (*N*)^3^, the semidilute regime, self-avoiding interactions are screened at lengths larger than the correlation length or “blob” size *ξ* = *aφ*^−3/4^, that is set by the monomer volume fraction *φ* = *kNa*^3^/*V* [1]. This scaling remains valid till the monomers fill the entire space, *φ* ≈ 1, the melt regime. The average volume fraction of the confined genome is in the semidilute regime [see Table S1], where the blob size is typically in the hundreds-of-nanometers (nm) range.

The chromatin blob size inside the brush, however, is smaller than the confinement-induced limit, because of the higher volume fraction driven by compaction. Loop extrusion-generated tension, *f* (in the pN-range) establishes a smaller blob size: ~ *k*_*B*_*T*/*f* (tens-of-nm range; recall, *k*_*B*_*T* ≈ 4 pN-nm), and drives a transition to a compact state where the local environment approaches the melt regime. Note that this transition is fundamentally different from that arising under “bad solvent” conditions, i.e, where instead of self-avoidance, inter-monomer contact is energetically favored. Unlike bad solvents, loop-extrusion, due to its topological nature, drives *only the monomers within a chromosome* towards a melt. Loop extrusion acts against chromatin self-avoidance and entropic mixing to preferentially *demix* chromatin associated with different chromosomes, and thus acts as an effective thermodynamic force driving segregation or individualization of chromosomes. This gives rise to an effective *chromosome polymer* that is made up of brush segments. In the following we analyze a homogeneous cylindrical brush, and use blobs made up of this shorter and stiffer brush-polymer to compute nearby contacts between inter-chromosomal segments as a measure of entanglement [7].

An important assumption in the above mentioned effective-polymer renormalization is that of *topology fluctuations*, ie., Topo II facilitates random strand passage. We treat chromatin as self-avoiding with the caveat that at long timescales corresponding to large structural reorganization of chromosomes, the effective polymers behave as “phantom” chains [2]. This means at short time scales of equilibration of blobs, there are topological constraints, however, when a certain constraint persist over time scales relevant to Topo dynamics (~ 1 strand passage per second), it is released with a ±1 linking number change. The relevant measure of genome entanglement in a transition from interphase to mitosis is the inter-chromosome linking number change. We find that under topology fluctuations, the distribution of inter-chromosome linking number is effectively controlled by lengthwise compaction. Lower compaction in interphase leads has a wider distribution, which becomes narrower under higher level of lengthwise compaction, driving Topo II-mediated unlinking or decatenation.

### A. Cylindrical polymer-brush chromosomes

We model chromosomes as a succession of chromatin loops connected by a chromatin backbone [Fig. 1(b)]. The resulting cylindrical bottlebrush polymer is characterized by three independent structural parameters: *loop size n*, the average number of nucleosome monomers per loop; *backbone size m*, the average number of nucleosome monomers between adjacent loop anchors; and *loop valency α*, the degree of subdivision (branching) of larger loops into smaller ones. A loop of size *n* with a valency *α* indicates there are *α* subloops each of size *n/α* associated with the same anchoring location [Fig. 1(b)].

When the backbone is comparable to the loops (*n* ≲ *m*), adjacent loops overlap only weakly, and the brush approaches the “random coil” limit [Eq. A1]. This corresponds to the average semidilute solution, where the monomer density inside and outside a chromosome are similar. Alternately, when *m* « *n*, adjacent loops strongly overlap and the resulting structure, resembling a “polymer brush” is significantly more compact, in addition to being stiffer.

A key parameter for the brush is the *interloop distance d*, the spatial distance between adjacent loop anchors: *d* is the steady-state end-to-end extension of the backbone segment of *m* monomers. A polymer brush morphology of overlapping adjacent loop-bristles requires *d* < *R*_*F*_ (*n*) and *d* > *R*_*F*_(*m*). For a fixed *m* and *n*, the value of *d* is set by a balance between the *osmotic repulsion* among adjacent loops that drives an increase in *d* (*f* ≈ *k*_*B*_*Ta*^5/8^*n*^3/8^*d*^−13/8^), and the *elastic restoring force* of the backbone polymer which favors a decrease in *d* (*f* ≈ *k*_*B*_*Ta*^−5/2^*m*^−3/2^*d*^3/2^) [Appendix]. This force balance furnishes: *d* = *an*^3/25^*m*^12/25^, that is valid until the stretching force reaches a critical value: *f*_*_ = *k*_*B*_*T/a* ≈ 0.4 pN, corresponding to a completely stretched backbone (*d* ~ *ma*). This transition occurs when the backbone size is a small fraction of the loop size, *m*_*_ = *n*^3/13^ [39].

As is typical for a cylindrical polymer brush, the monomer volume fraction decreases radially outwards causing an osmotic pressure gradient that radially stretches the loopsand establishes a long thermal bending persistence length reflecting a stiffening response for a brush with closely spaced loops [Appendix] [34, 39–41].

The average monomer volume fraction inside the brush is higher for lower interloop distance and higher loop branching: *ϕ* ~ *α*^2/3^*d*^−2/3^. This generates a region along the backbone of the brush, we refer to as the “core”, that features dense packing of the monomers (*ϕ*_*core*_ ≈ 1) and a high osmotic pressure ∼ *k*_*B*_*T/a*^3^ [Fig. 1(c-d)]. The width of the core is proportional to the loop valency and inversely proportional to the interloop distance: *r*_0_ ≈ *αa*^2^*/d* [Appendix]. This relation, derived from the geometric condition that the surface of the core form a surface saturated by “grafted” loops, indicates that more closely packed loops and loops with higher branching generate a thicker core. We note that while the existence of maximal-density core has been long discussed in connection with spherical polymer “micelles” [42], the cylindrical polymer brush literature has not recognized this possibility, which appears to be key to understanding chromosome folding.

### B. Brush chromosomes as an effective polymer

We consider the brush chromosomes as a thicker and shorter “noodle”-like polymer constituted of the underlying chromatin. The renormalized brush contour length *L′*is shorter than the chromatin contour *L* = *Na*. The renormalized thickness *R* is larger than chromatin thickness *a*, and shows a monotonic increase with loop size. However, *L′*is non-monotonic with loop size *n*:

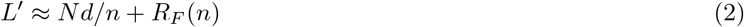

The first term corresponds to the cylindrical part of the brush where there are *N/n* loops, each contributing *d* ∼ *n*^3/25^*m*^12/25^ to the axial length *L′*–constituting a net contribution that decreases with the loop size. The second term is from the loops at the ends of the cylinder that are less confined than their counterparts in the middle of the brush. These hemispherical ends of the chromosome have radii on the order of the chromosome thickness and contain a large amount of chromatin (e.g., megabases for larger human chromosomes), and are typically much larger than telomeres. As the average loop size increases, the contribution of the “end loops” to the axial length increases until one has a “star” polymer which is all ends and no axis, with size of a Flory polymer of *n* monomers [Eq. A1]. We note that Eq. 2 corresponds to the brush limit of the chromosomes where *n* » *m*, or *n* + *m* ≈ *n*.

Minimization of the renormalized contour length gives an *optimal loop* size:

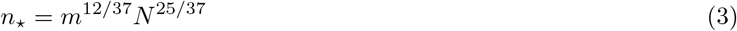

that maximizes compaction. A renormalized chromosome with loops larger than the optimal size begins to resemble a “star” polymer or a micelle, whereas, loops smaller than the optimal size makes the structure a thin brush with a long axial length. Both of these non-optimal possibilities are less compact than the *optimal loop* brush.

### C. Entanglement between chromosomes in a confined volume

We quantify entanglement between chromosomes using their mean-squared inter-chromosome linking (catenation) number 〈Ca^2^〉, which measures the width of the catenation number distribution. 〈Ca^2^〉 is readily computed from the number of near (polymer segment-scale) encounters between the brush chromosomes [6] (the ‘catenation number’ Ca is used in the DNA topology field to denote Gaussian linking number of distinct duplex DNA molecules). Given freely fluctuating topology, each close encounter between chromosomes contributes ~ ±1 to the inter-chromosome linking number. Since Topo II is a locally acting enzyme that cannot sense the global linking between any two chromosomes, the only means chromosomes may disentangle in presence of Topo II-mediated strand-passage activity is via elimination of inter-chromosomal contacts, which may be achieved by driving higher compaction.

The number of close encounters or collisions between chromosome segments, that determines the level of entanglement, scales linearly with the number of polymer segments: 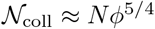 [6] [Appendix]. As previously shown in Ref. [6], the contribution of this blob-collision number to inter-chromosomal topology is obtained using 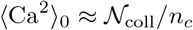, where the proportionality constant *n*_*c*_ ≈ 100 (determined numerically [6, 43]) is the number of contacts required to have a ± 1 contribution to catenation (*n*_*c*_ is a scale similar to the “entanglement length” familiar from polymer physics [1]).

The number of collisions between segments of the effective brush polymer, 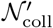 is much lower than that for chromatin 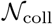 due to the smaller number of statistical segments of a polymer brush *N′* ≈ *L′*/*R* « *N*. This leads to an inter-chromosomal entanglement level for the brush state (〈Ca^2^〉) that can be controlled via the compaction state of the chromosomes, and is significantly lower than that for unfolded chromatin (〈Ca^2^〉_0_):

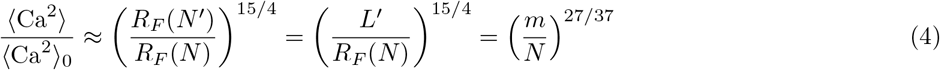

where the last two equalities follow for optimal brushes (*N′* ~ 1). Since *m* » *N* there is a strong suppression of the entanglement level in a solution of optimal brushes, compared to a semidilute solution of linear chromatin chains (for calculation details see SI).

We have used a flexible self-avoiding polymer approach for describing chromatin (with nucleosome-scale monomers); we also employ this description for bacterial “chromatin” (DNA covered by nucleoid-associated proteins or NAPs), as well as for the conformational statistics of the “brush” polymers formed by loop extrusion. In the bacterial chromatin and brush-polymer analysis, we note that for situations where the polymer segments are much longer than their width, the weak excluded volume between monomers might oblige one to follow the approach of Ref. [44], which considers the situation of long, thin polymer segments. For this limit, one has a concentration range in which excluded volume is too weak at short distances to generate self-avoiding-walk statistics, and where one should use marginal or “theta” solvent conditions. For the situations considered in this paper, we have found that effects of segment shape are not critical to the results, so we follow the simpler self-avoiding polymer formulation; discussion of segment-aspect-ratio effects can be found in the Appendix.

## II. RESULTS

### A. Optimal loops maintain a basal level of chromosome compaction and suppress inter-chromosomal entanglements

Optimal loop size minimizes chromosome axial length *L′*[Eq. 2]. The existence of the optimal state arises from the fact that when the loops are small, there are a large number of them, each contributing about one interloop distance *d* to *L′*. On the other hand, if the loops become so large that the equilibrium unperturbed size of the loops at the end of the cylindrical brush begins to dominate, *L′*is again large. A loop size between these limits minimizes *L′*. Figure 2 shows the optimal subloop size, which is obtained by dividing the optimal size by the valency or the number of loop branches.

**FIG. 2.**
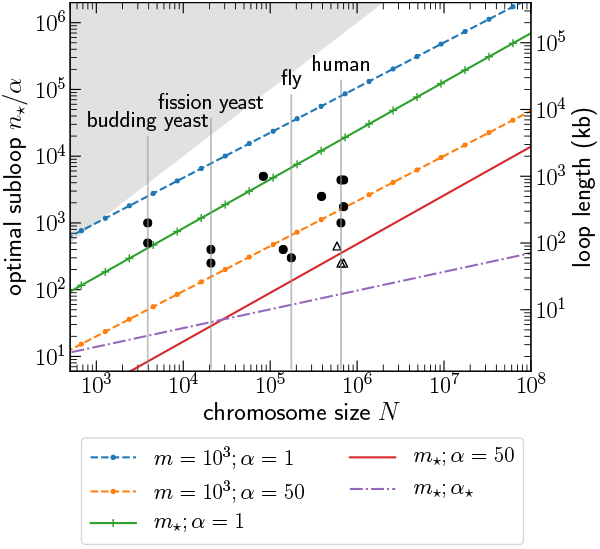
Optimal loops of size *n*_*_ minimize axial length of the brush chromosome and suppress chromosomal entanglements. Loop size divided by valency *α* (number of branches), gives the size of subloops, plotted versus the total number of chromosomal units *N*. Curves correspond to different number of monomers in the backbone segment between two consecutive loops *m* and valency *α*. Optimal loops are larger for larger *m*. A fully stretched backbone, owing to the tension generated from overlapping adjacent loops, corresponds to *m* = *m*_*_ [Table I], the minimum size of the backbone for an optimal loop. Higher valency *α* leads to smaller subloops due to increased loop branching and increases the density of the monomers in the interior of the brush. Valency *α*_*_ scales with *N*, and corresponds to a phenomenological value for metaphase chromosomes [Table I]. Gray-shaded region is inaccessible, as it corresponds to a loop size greater than the chromosome length. Filled circles are experimental data for “loop domains” in the interphase genome obtained from chromosome contact or Hi-C maps [28–30, 32, 33]; open triangles denote loops identified from electron-microscope images of metaphase chromosomes [9, 10]. The data indicate that an entanglement-suppressing optimal loop size is maintained throughout the cell cycle: during interphase, the loops have a lower valency and are less compact, whereas during mitosis, the loops are heavily branched, leading to smaller subloops. The experimental data also respect the physical limits imposed by our tandem-loop model. The *y*-axis on the right shows the loop lengths in kilobase pair (kb) units.

**TABLE I.**
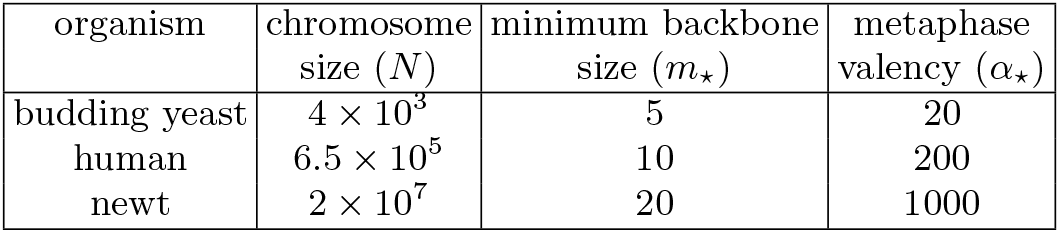
Example values of the minimum backbone size: *m*_*_ = *N* ^15/89^, and metaphase valency: *α*_*_ = (0.5)*N* ^40/89^

In terms of inter-loop-anchor “backbone” segment size *m*, optimal loops are larger for a brush with larger *m* (Fig. 2). Since, to attain the same brush axial length and stretching tension along the backbone, a configuration with longer backbone segments requires larger loops. Increasing *m* eventually leads to non-overlapping adjacent loops, the case for a random-coil polymer. Making the backbone small leads to the limiting size *m*_*_, where the tension from overlap of closely anchored loops completely stretches the backbone (*d* ~ *m*_*_ ~ *N*^15/89^) [Table I]. The optimal brush with a fully stretched backbone has the minimum axial length *L′*, associated with an axial stretching tension *f*_*_ = *k*_*B*_*T/a* ≈ 0.4 pN. Compaction with sub-pN stretching forces is not likely to disrupt the nucleosomes or the integrity of the chromatin fiber, an important physical constraint on genome folding.

The different lines in Fig. 2 correspond to values of *m* and *α* that represent different overall conformations of brush chromosomes. The blue dashed line (*m* = 10^3^ and *α* = 1) corresponds to a “sparsely grafted” configuration of monovalent loops, featuring minimal overlap between adjacent loops. While, the orange dashed line (*m* = 10^3^ and *α* = 50) corresponds to a sparse configuration of *branched* loops. The green solid line (*m* = *m*_*_ and *α* = 1) corresponds to a “dense” regime of brush where adjacent monovalent loops are closely grafted, forcing strong inter-loop repulsion and a stiff response to bending. The red solid line (*m* = *m*_*_ and *α* = 50) is for a dense brush with higher degree of loop branching. Finally, the purple dot-dashed line (*m* = *m*_*_ and *α* = *α*_*_) corresponds to a dense, stiff brush with highly branched loops – a model for compact metaphase chromosomes; the degree of branching *α*_*_, which scales with *N* (see below and Table I), is determined from the elastic modulus of metaphase chromosomes [45].

Hi-C experiments studying the ensemble-average conformation of interphase genome of various species find a characteristic loop size [29, 31–33] (Fig. 2, filled circles). Fig. 2 suggests that these *interphase* chromosome loops maintain a brush-like structure of the chromosome, driving a level of compaction quantitatively similar to optimal loops. Loop sizes obtained from electron-microscope images of metaphase chromosomes [9, 10] are somewhat smaller than interphase loops (Fig. 2, open triangles). Increasing interphase loop valency is a conceivable way to drive mitotic chromosome compaction, but Hi-C experiments do not indicate any sequence-specificity of loops during mitosis [25, 31], suggesting a major refolding of the genome that is stochastic in nature.

### B. Chromosomes can be completely disentangled via compacting optimal loops

Optimal brush chromosomes are semiflexible polymers, because the thermal persistence length of optimal chromosomes are comparable to its axial length. The long persistence length is a result of compaction tension along the backbone, which makes the brush stiff to bending fluctuations. This leads to a significantly lower number of statistical segments of the optimal brush than that of the constituting chromatin (*N′* « *N*), causing a lower level of entanglement between brush chromosomes [Eq. 4].

The equilibrated-topology entanglement level 〈Ca^2^〉[Eq. 4] may be estimated as proportional to the number of inter-chromosomal contacts per chromosome; each contact represents a point where a crossing between two genomic segments can be reversed in sign without significant perturbation of chromosome conformation [7]. Unfolded chromosomes generally show a high degree of inter-chromosomal entanglement [Fig. 3(a)], since the scale of nuclear confinement is smaller than the unperturbed equilibrium random-coil size of unlooped chromosomes [Eq. A1] [1].

**FIG. 3.**
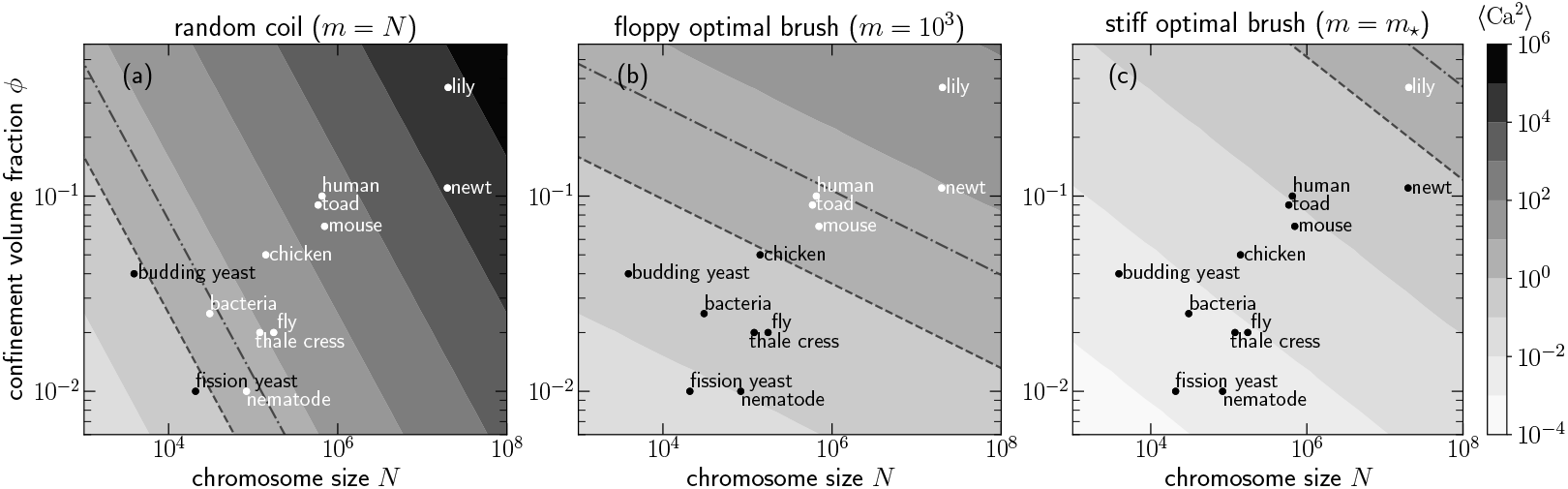
Removal of inter-chromosomal entanglements by establishing an optimal brush configuration via loop extrusion in the presence of topology fluctuations. A cell with a given nucleus volume, number of chromosomes, and an average chromosome size *N* occupies a specific position in the above (*N, φ*) diagram, where *φ* is the net volume fraction of all the chromosomes within nuclear confinement. Shading indicates level of entanglement 〈Ca^2^〉[Eq. 4], where a lighter (darker) shade depicts a lower (higher) entanglement or inter-chromosomal contacts per chromosome (see legend to the right). Dashed line shows 〈Ca^2^〉 = 1 in the nucleus: above this line cellular chromosomes are entangled (〈Ca^2^〉 *>* 1), while a cell lying below the dashed line has a disentangled genome (〈Ca^2^〉*<* 1). Dot-dashed line shows 〈Ca^2^〉= 1 line after nuclear envelope breakdown (NEB), where the confinement volume is increased 3-fold over that of the nucleus; NEB mildly aids chromosome disentanglement. (a) Chromosomes considered as random coils of self-avoiding polymers exhibit a significant degree of entanglement (〈Ca^2^〉_0_ ~ *Nφ*^5/4^), that increases with the chromosome size. Notably, yeast chromosomes are essentially disentangled in the random coil state. (b-c) Chromosomes modeled as polymer brushes are more compact and as a result are less entangled. Entanglements between chromosomes can be reduced by organizing the chromosomes into a tandem array of optimal loops, which have a minimal end-to-end extension for a given backbone segment size *m*. Contour plots show that steady-state optimal brush chromosomes have a lower entanglement level than the random coil state. Further removal of entanglements is possible by stiffening the optimal brush via stretching and shortening the backbone. Interphase chromosomes have larger loops separated by long backbone segments, making them less compact, floppy brushes that are also less entangled than a random coil. Loop extrusion mechanism can modulate the brush configuration to generate an order *k*_*B*_*T*/*a* ≈ 0.4 pN tension that completely stretches the backbone (*m* = *m*_*_), ultimately leading to stiffening and complete disentanglement of chromosomes.

Higher confinement volume upon nuclear envelope breakdown (NEB) is a mild effect, and by itself, does not strongly drive segregation of chromosomes [Fig. 3(a)]. Interestingly, nematodes and yeast have essentially disentangled genomes even in a random-coil state, indicating that segregation of chromosomes is a less pressing concern for these organisms compared to, e.g., mammals, whose chromosomes can get highly entangled. Some lower eukarya (e.g., budding yeast) are known to *not* have NEB during mitosis, i.e., “closed” mitosis [46]; the essentially disentangled genome inside the nucleus of these organisms may have contributed to this evolutionary outcome.

The entanglement level is much lower (for given confinement volume fraction and chromosome size) when the chromosomes are organized as a tandem array of optimal loops [Fig. 3(b)], highlighting the importance of loops during interphase. Figure 3(b) is plotted for *m* = 10^3^, a regime in which the optimal brush is relatively “floppy”, i.e., the backbone size is such that adjacent loops moderately overlap (*m*_*_ < m < *n*_*_). While a smaller backbone size (*m* = *m*_*_, Fig. 3(c)) corresponds to a stiffer and shorter brush.

Figure 3(a) shows the maximum possible entanglement between chromosomes for a given confinement volume fraction and average (linear) chromosome size: due to interphase looping, chromosomes are never strongly entangled. Reducing backbone size (lower *m*) further lowers the level of inter-chromosomal entanglements [Fig. 3(c)]. Stretching and shortening the backbone leads to a stiffening response of the optimal brush that drives the disentangled state. Given typical nuclear volume fractions, stretching the backbone of optimal brush chromosomes (force 1 ≈ pN) completely disentangles them.

### C. Compact chromosomes have a dense axial core featuring closely packed monomers that impart mechanical rigidity

Compaction of the optimal brush chromosomes can be controlled via two physical processes: one, shortening the backbone (lower *m*); and two, increasing the valency of the optimal loops (higher *α*). Both these processes increase the monomer concentration in the interior of the brush, leading to an overall compaction; however, the signatures of these processes on the compacted structure are different: shortening the backbone compacts the axial length [Eq. 2], while, increasing the average valency of optimal loops compacts the lateral dimension or thickness of the chromosomes: *R* ~ *α*^−1/2^.

Higher monomer concentration inside the brush leads to a densely packed core where the monomer volume fraction is near maximal (≈1), and consequently, the core has a high osmotic pressure ≈*k_B_T/a*^3^ ≈4 kilopascals (kPa). This contrasts with the much smaller bulk modulus for an ordinary (uncrosslinked) semidilute polymer solution ≈10 Pa, due to a lower volume fraction ≈0.1. The elastic modulus of brush chromosomes shows a strong dependence on the average loop valency: *E ~ α*^9/4^. We use the experimental value of metaphase chromosome elastic modulus *E*_meta_ ≈ 1 kPa [45, 47, 48] to determine the *metaphase loop valency* for an optimal brush with fully stretched backbone: *α_*_ cN* ^40/89^. Here *c* = (*E*_meta_*a*^3^*/k_B_T*)^4/9^ ≈ 0.5 is an order-unity constant [Table I].

Figure 4(a-c) show the structural aspects of the brush chromosomes for various *m* and *α*, where the legends are identical to Fig. 2. The axial contour length in our model does not depend on loop subdivision, so Fig. 4(a) shows only two curves corresponding to different values of *m*. This indicates that there is not a large decrease in axial contour length during the later (prometaphase) stages of mitotic compaction where loops are subdivided (*m* increased) [25]. The thickness, however, undergoes significant compaction upon subdivision of loops [Fig. 4(b)].

**FIG. 4.**
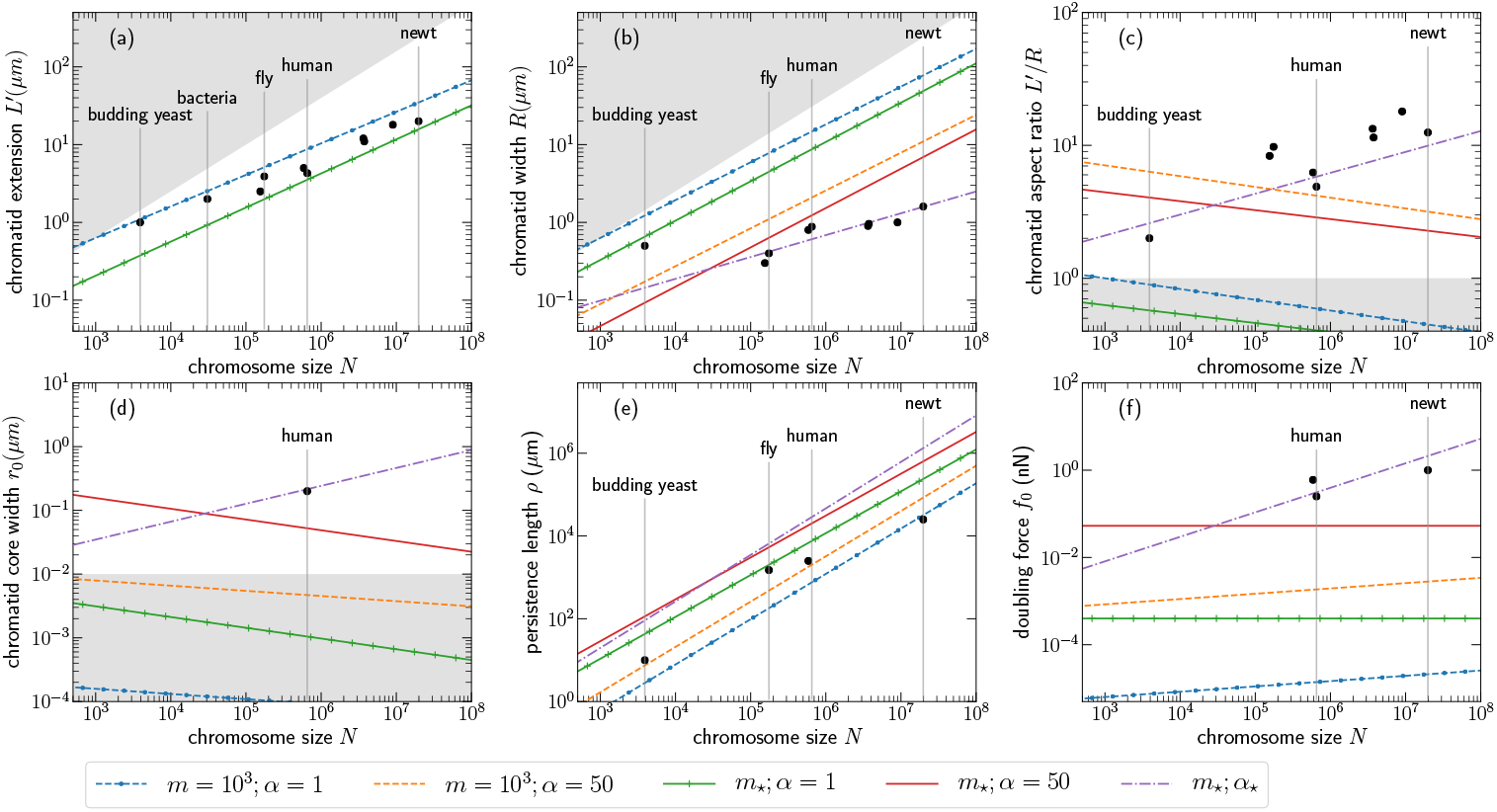
Structure and mechanics of compact mitotic chromosomes result from the densely-packed chromatin core. Filled circles show experimental values for metaphase chromosomes [see SI]. For the bacterial case, filled circle shows the dimension of a cellular chromosome which compares with the cellular dimensions. Lines correspond to the cases described for Fig. 2. (a) Average chromatid axial length *L′*, (b) chromatid width *R*, and (c) length-to-width aspect ratio (*L′*/*R*) show that the change in width is stronger upon stiffening the brush via shortening the backbone (lower *m*) and increasing the loop valency (higher *α*). Larger valency values promote a higher monomer concentration in the interior of the brush and generate a core where the monomers are densely packed. The core enhances the mechanical rigidity of the chromosomes; the width of the core *r*_0_, which is proportional to the average loop valency, is plotted in (d); the lines lying in the shaded region in (d) do not have a core, since a minimal core must be at least one monomer thick. The experimental data for human chromosome core corresponds to the thickness of the axial region where condensin-colocalizes in metaphase [23]. The thermal persistence length of the brush *ρ* (e), and the force associated with stretching the chromosome to twice its native length or the doubling force *f*_0_ (e), show increased stiffness and mechanical rigidity for smaller backbone and higher valency. The valency *α*_*_, which is required to reproduce the observed metaphase-chromosome elastic modulus ≈1 kPa and is ≈40% of the maximum allowed valency of an optimal loop, agrees well with metaphase chromosome size data. This indicates that the dense core resulting from backbone stretching and branching of chromosomal loops, achievable via a loop extrusion mechanism, can underlie the emergent mechanical rigidity and compactness of metaphase chromosomes. Shaded region in (a) and (b) correspond to chromosome for which the backbone is comparable to the loop size leading to no overlap between adjacent loops, which approaches the random coil limit of the polymer brush: *L′* ≈ *R*_*F*_ (*N*) and *R* ≈ *R*_*F*_ (*N*). This suggests bacterial chromosomes are only minimally compacted in the axial dimension. Also, note that backbone segments with *m* = 1000 monomers correspond to overlapping adjacent optimal loops for all relevant chromosome sizes.

Comparison with metaphase data suggests that metaphase chromosomes have a sufficiently stretched backbone (*m* = *m_*_*) supporting highly branched loops (*α* » 1). The core for metaphase chromosomes of higher eukarya is expected to be ≈100 nm thick [Fig. 4(d)]. The thermal bending persistence length, and the doubling force (force required to stretch chromosomes to twice its native length: an intensive quantity) are measures of mechanical rigidity, and show strengthening of the brush in metaphase due to formation of the thick core [Fig. 4(e-f)] [Appendix]. Metaphase doubling force originates primarily from the core, however, this force is not equivalent to the polymer backbone stretching force that increases interloop distance *d*. This force is a product of the high osmotic pressure generated in the core from dense packing, which possibly depends heavily on chromatin crosslinking. Additionally, local adhesion among the core monomers may also provide stabilization.

### D. Bacterial chromosomes

Bacteria, unlike eukaryotes, do not have nucleosomes; instead, bacterial DNA inside cells is coated with a variety of nucleoid-associated proteins (NAPs), e.g., HU, H-NS and IHF [49]. Like eukaryotes, bacteria possess SMC proteins (in *E. coli*, MukBEF [50]) and bacterial versions of eukaryote TopoII. We treat bacterial chromosomes as self-avoiding polymers, with cylindrical monomer units of length *a* ≈ 50 nm (comparable to the persistence length of naked DNA [51]) and width *b* ≈ 5 nm, corresponding to the thickness of protein-bound DNA segments (NAPs can reduce the persistence length but this is not crucial here). This gives a monomer aspect ratio constant *a/b* ≈ 10, that shows up in the ratio between volume and excluded volume of a monomer. The macroscopic (renormalized) lengths, *L′* and *R*, scale weakly with the aspect ratio, making the structure optimization and entanglement discussions for spherical monomers applicable to bacterial chromosomes.

The bacterial genome has the potential to be appreciably entangled due to its substantial confinement [Fig. 3(a)], highlighting the need to drive compaction in order to segregate multiple copies. However, only mild compaction compared to higher eukaryotes is required [Fig. 3(b)]. Bacterial DNA is also subject to a global supercoiling pressure, by virtue of DNA gyrase, a motor-like enzyme that maintains bacterial DNA in a supercoiled condition. Supercoiling may play an important role in driving compaction and maintaining an optimal brush conformation of bacterial chromosomes. The restoring force of a plectonemic domain for physiological levels of bacterial supercoiling ≈0.5 pN [52], can generate the stretching tension along the backbone connecting plectonemic domains, necessary to drive compaction and segregation of bacterial chromosomes. This is in accord with models of bacterial chromosome organization into territories driven by DNA supercoiling [53].

For elongated bacterial cells that are asymmetric (e.g., *E. coli* and *C. crescentus*), there is expected to be an entropic-segregation pressure gradient along the long axis of the cell. This may aid chromosome segregation by pushing the two sister chromosomes to opposite poles of the long axis [54]. The cylindrical brush structure of chromosomes, driven by loop extrusion, enhances this pressure gradient, and provides an active mechanism to control segregation. Other mechanisms may work in parallel: some bacterial species possess chromosome tethering mechanisms that may aid in driving axial segregation [55].

The optimal loop result for the *E. coli* nucleoid of axial length 2 *µ*m corresponds to a floppy brush (blue dashed line in Fig. 4(a)), which permits full segregation but with enough flexibility to allow the nucleoid to be folded and moved around inside the cell. The bacterial nucleoid is heavily confined, evident from the large expansion (3 to 10 fold linear dimension) of the bacterial chromosome following cell lysis. Bacterial nucleoids removed from cells behave as polymer networks of roughly 10 *−* 20 *µ*m maximum extension, [55, 56], consistent with the maximum extension of a loosely compacted brush of axial length *≈* 2 *µ*m.

## III. DISCUSSION

We used a polymer model to describe cellular chromosomes in confinement and showed that a chromosomes structure characterized by an array of connected chromatin/DNA loops, in presence of topology fluctuations via random strand passage by Topo II, exhibits lower inter-chromosomal entanglement than the corresponding unfolded, linear-polymer configuration. We found that folding a chromosome in loops of a chromosome-size-dependent length –the “optimal” loop length– simultaneously maximizes compaction and minimizes inter-chromosomal entanglements [Figs. 2 and 3].

These optimal loops (or, subloops formed upon branching) are comparable to experimentally observed loops through the cell cycle, suggesting a role of chromosomal loops in suppressing entanglements, both during interphase and mitosis. These loops keep the chromosomes territorialized throughout the cell cycle, tightly regulating interchromosome entanglement. Larger loops and a longer backbone allow a controlled level of entanglement in interphase, which is likely essential for gene expression. While the positioning of these loops may affect transcriptional regulation, correspondence of their sizes to that of entanglement-suppressing optimal loops may be linked with the evolutionary selection of chromosome architecture.

On the other hand, compact mitotic chromosomes have short, stretched backbones with heavily branched loops, that efficiently remove all the entanglements between chromosomes leading to their segregation [Fig. 3]. Mitotic chromosomes have a densely-packed core along their cylindrical axes. The high density of monomers in the core is responsible for the emergent mechanical stiffness of chromosomes during mitosis. The predicted size of mitotic subloops required to reproduce the experimentally observed chromosome stiffness is smaller than the subloops observed in Electron-microscope images [Fig. 2]. A larger subloop in our model will lead to a lower chromosome stiffness moduli, however, the mechanical rigidity may be compensated by additional crosslinking of chromatin inside the chromosome brush, which we ignore in our simple approach.

### A. Topo II-driven topology fluctuations allow segregation and disentanglement via compaction

Topo IIs allow passage of chromatin segments through one another, permitting chromosome topology to fluctuate [2]. However, Topo IIs cannot disentangle chromosomes by themselves [57–60], since they are unable to directly sense global chromosome topology. By allowing topology fluctuations, Topo II can maintain topological equilibrium, allowing disentanglement to occur gradually as lengthwise compaction proceeds. Our theory shows that the entanglement level tracks the chromosome architecture and its manipulation by loop extrusion (see below). Further work on the time evolution of entanglement release is a next step: DNA tension at interlocks and effective viscosity generated by entanglements are likely to be important.

We have assumed the null hypothesis of *random* strand passage by Topo II; a synergistic mechanism where Topo II directly interacts with SMC complexes has been suggested to drive more efficient disentanglement [5]. Intra-chromosomal topology, such as chromatin knots, have also been shown to be suppressed by a combined action of SMC and Topo II [4]. In our model, loop-extrusion folding of chromatin by SMCs provides a local free-energy gradient pushing Topo II to resolve inter-chromosome catenation, thus also coupling SMC activity to topology changes mediate by Topo II.

### B. Loop extrusion can control the optimal brush structure

Loop extrusion has emerged as a vital mechanism underlying organization of chromosomes. SMC complexes (cohesin and condesnsins) can exert pN forces and are the prime candidates for driving the loop extrusion activity [26, 27, 61–64]. In our model, extrusion of loops generates interloop repulsion that stretches the backbone; thus, loop extrusion may effectively control *n*, *m*, and *α* to drive chromosome compaction, and consequently, disentanglement in presence of Topo II. The ubiquitous presence of DNA-bound SMC complexes in all cells can ensure a steady-state stretching of backbone segments, important for maintaining a semiflexible chromosome brush, and a low entanglement level between confined chromosomes throughout the cell cycle.

The Rouse equilibration time of a typical optimal loop (100-1000 monomers) is less than a second, with topology changes requiring roughly a second per strand passage cycle. The relevant time scale associated with loop extrusion activity is not clear, individual SMC motors have been observed to translocate on DNA at ~ 1 kb/s [27]. However, any large scale structural reorganization involving Mbp-sized chromosome loops is expected to occur over at least many minutes (i.e., in mammals, on the order of 10 traversals of ≈ 100 kb loops of chromatin by SMCs moving at kb/s speeds, or roughly ≈ 10^3^ s, similar to the duration of mammalian prophase), much longer than the Rouse time of the loops. This allows us to use static scaling laws that are governed by the elasticity of chromatin. Analysis of the dynamics during the reorganization process may be possible by combining relaxational polymer dynamics, topology change by Topo II, and estimates of non-equilibrium active forces generated by SMC complexes.

### C. Chromosome loop organization during interphase

The size of the interphase loops and their positioning depends on proper functioning of the loop-extruding SMC complexes. Destabilization of interphase loops (possibly via inhibition of SMC activity) is expected to disrupt the optimal structure, resulting in higher chromosomal overlap, i.e., weakening of chromosomal territories, as has been observed [22]. We also expect a concomitant increase in the inter-chromosomal entanglements for less territorial chromosomes.

Compartmentalization of the genome into early- and late-replicating domains (respectively, eu- and heterochromatin), may be relevant to maintaining low genome entanglement during DNA replication. Replication is expected to cause disassembly of loops, especially when replicating the loop anchoring regions, likely leading to an overall increase in entanglement between chromosomes. A controlled disassembly of loop organization via compartmental replication (and possibly, restoration of loops immediately following replication) restricts large-scale entropic mixing of chromosomes during S-phase.

Various non-SMC complexes, such as CTCF proteins are integral to chromosome architecture and are possibly relevant for the optimal structure. CTCF proteins are known to stabilize certain sequence-specific loops that play a role in gene regulation [32], some of these loops have also been reported to remain stably bound throughout the cell cycle [65, 66]. Such structural template of connected loops is capable of storing heritable gene expression patterns while simultaneously minimizing chromosomal entanglements.

DNA twisting or supercoiling pressure is another important aspect of chromatin, especially during interphase, when the DNA is transcribed and replicated. Supercoiling flux, e.g., at the loop anchors may regulate the level of loop compaction, and stabilize certain loops [53].

### D. Chromosome structural rigidity and topological disentanglement during mitosis

Loop extrusion activity by SMC complexes maximizes compaction by minimizing the axial length of brush chromosomes. Condensin II is a likely candidate that drives the prophase compaction, suggesting an important role of condensin II in determining the axial length of chromosomes, in accord with the observation of an increase in the axial length for condensin II-depleted chromosomes [17, 23, 67–69]. Note, due to the optimal loop architecture of chromosomes in interphase, we do not expect the axial length to significantly change during the course of the cell cycle – which is central to our conclusion that inter-chromosomal entanglements are suppressed by SMC activity during the cell cycle.

The other SMC complex, cohesin is known to hold the sister chromatids during mitosis, however their role in prophase compaction of chromosomes, if any, is not clear. If indeed condensin II is solely driving prophase compaction and segregation, inactivation of cohesin activity in prophase will lead to a factor-of-two increase in the number of chromosomes, which is *not* predicted to be crucially detrimental to their segregation [Fig. 3(c)].

The radial dimension of cylindrical chromosomes is strongly compacted by loop division, which establishes a dense core along the axes [Fig. 4]. Higher valencies corresponding to branched loops in metaphase may occur for eukaryote chromosomes via binding of condensin I after nuclear envelope breakdown [25]. The observation that condensin I-depleted chromosomes have a thicker diameter and a lower stiffness [17, 68, 70], supports the notion that these proteins branch loops established by condensin II during prophase, to generate rigid, rod-like metaphase chromosomes.

SMC activity is crucial for the dense core in metaphase chromosomes. The high osmotic pressure from close packing of nucleosomes in the core is possibly stabilized by additional mechanisms such as nucleosome-nucleosome attraction in mitosis [71], and a higher concentration of chromatin-crosslinking proteins (including Topo II) inside the axial core. A recent proposal, supported by Hi-C studies, posits that SMC complexes drive a helical brush axis during metaphase [25]. Such phenomenology may be studied within the framework of our model, and is left for future discourse.

### E. Segregation of sister chromatids from osmotic repulsion between the axial cores

Loop extrusion activity on the newly replicated catenated sister chromosomes leads to two brushes that are intertwined near their backbones; the overlapping loops generate a repulsive interchromatid force. The net repulsive force follows from the total osmotic pressure per cross-sectional area of overlap between the two sister chromatids, giving a force acting on each loop of *f*_*rep*_ ≈ 10 pN for monovalent loops, and increases strongly for higher loop valencies *f*_*rep*_ ~ *α*^1/2^ [Appendix]. This repulsion drives strand passages by Topo II that will lead to physical segregation of sister-chromosome brushes.

Importantly, an indiscriminate increase of loop valency and crosslinking while the sister-chromatid backbones are still heavily entangled will lead to fusion of the sister chromosomes into a common core, hindering their segregation. This predicts heavily entangled sister chromosomes if condensin I is active inside the nucleus *during prophase*. Initiation of condensin II compaction and sister chromosome segregation immediately following replication [72, 73], is crucial for timely removal of entanglements *before* establishing a thick core (i.e., before condensin I activity). Once the backbones are disentangled, removal of residual entanglements between sister chromatid loops are facilitated by core formation, since the repulsive force between chromatid backbones is higher for a thicker core. Our model rationalizes the sharp, disjoint compartmentalization of condensin II and condensin I in terms of a kinetic process: establish packed loop arrays first (condensin II) and *then* generate a dense core by loop valency increase (condensin I), and is in line with conclusions drawn from Hi-C analyses [25].

Fusion of the sister chromatid cores near the centromeric region is unavoidable (the sisters remain attached there until anaphase) High concentrations of Topo IIs and SMCs at the centromere during metaphase, resulting from the high DNA density, will generate strong repulsive forces between the dense and short centromeric loops possibly playing a crucial role in disentanglement of the (peri)centromeric regions [74, 75].

The smallest length scale in our analysis is the nucleosome/monomer size (≈ 10 nm), so we do not explicitly consider effects of electrostatic interactions which are of shorter range (the screening length is ≈ 1 nm under the 0.15 M univalent salt conditions found in the nucleus). Of course, local interactions generated by electrostatic interactions, especially due to divalent or multivalent charged species play a key role in chromosome organization e.g., local adhesion between nucleosomes or DNA-site-specific interactions, but for this paper we are concerned with larger-scale chromosome organization.

In conclusion, we have developed a steady-state polymer brush model for chromosomes, where the brush structure is primarily controlled via loop extrusion. Our major result is that the loop organization of the cellular genome is an entanglement-suppressing structure, explaining experimental observations. Lengthwise compaction of the brush results in chromosome segregation, establishing that loop extruders are capable of driving chromosome individualization and chromatid segregation.

## Acknowledgement

The authors acknowledge funding from NIH grants R01-GM105847, U54-CA193419 (CR-PS-OC) and U54-DK107980 (4DNucleome). We thank E. Banigan, A.D. Stephens, and R. Biggs for their helpful discussions.

## Appendix A: Scaling calculations: Semidilute solution of polymers with fluctuating topology

We adopt the notion of a polymer as a series of non-overlapping deGennes’ blobs [1], where the blob size or the correlation length scales inversely with the monomer volume fraction: *ξ* = *aϕ*^−3/4^. And the statistics of the polymer inside a blob is that of a self-avoiding walk or a Flory walk [1, 76]:

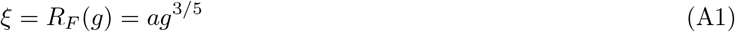

where *g* is the number of monomers of diameter *a* in each blob. Whereas, the chain as a string of blobs exhibits ideal or Gaussian polymer statistics. We have assumed good solvent conditions throughout the article, which is necessary to avoid collapse of all the chromosomes into one genomic globule. However, an adhesion scheme that –favors monomer-specific (epigenetic) contacts, is only locally operative, and *not* strong enough to unravel the overall loop organization– appears to reproduce experimental signatures [77, 78]. The framework of this paper can be extended to analytically study local adhesion and looping degrees of freedom.

Throughout this study we assume the topology to be fluctuating, which is a consequence of the presence of TopoII that allows passage of DNA segments through one another at a close-contact site. In such a fluctuating topology ensemble, the number of contacts between chromosome blobs is a measure of entanglement, and disentanglement is possible only when the number of inter-chain contacts is negligible.

### 1. Cylindrical polymer brush chromosomes

A cylindrical polymer brush has radially increasing blob size, since the net volume accessible to the loops is higher at a larger radial distance resulting in a radially decaying volume fraction.

#### a. Radial profile of monomer volume fraction

Following the arguments originally proposed by Daoud and Cotton [42] for the shpherical symmetry of star polymers, we consider a cylindrical shell of height *d*, inner radius *r*, and outer radius *r* + *ξ*(*r*). Since the blobs diffuse radially outward due to the monomer concentration gradient, and there is, on average, *α* blobs in the considered shell, where *α* is the loop valency. The volume fraction of monomers in the shell, *ϕ*(*r*), is equal to that inside a blob in that shell, leading to the following equations [39]:

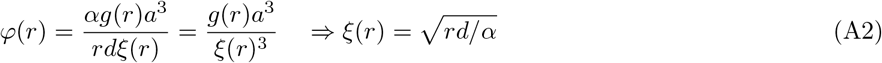

where *g*(*r*), the number of monomers per blob of size *ξ*(*r*) [(A1)].

#### b. Osmotic pressure exhibits radial decay

The osmotic pressure inside the brush Π, scales inversely with the correlation length [1]:

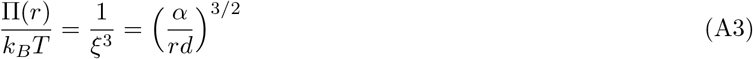

#### c. Loop extension

The radial extension of a loop, *R*, containing *n* monomers can be obtained from radial integration of the volume fraction *ϕ*(*r*) [34, 39, 41].

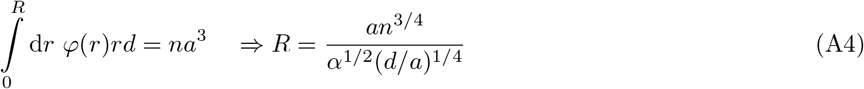

The scaling behavior *R* ~ *n*^3/4^ resembles a 2D-self avoiding walk, which is a consequence of the effective confinement of a loop in a slit-like geometry due to its neighbors [79].

#### d. Loop free energy

Free energy per loop is given by the number of blobs per loop, because each blob contributes ≈ 1 *k*_*B*_*T* [1]. Equivalently, the free energy per loop may also be obtained from integrating the total osmotic pressure in the cylindrical volume accessible to each loop, as follows [39, 41].

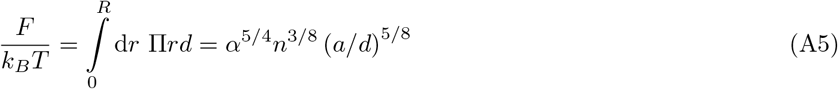

#### e. Tension along the backbone from loop overlap

Overlapping loops generates a higher osmotic pressure in the overlap volume that causes repulsion between adjacent loops and leads to a stretching tension along the backbone. We estimate the tension from the free energy per unit length of the backbone.

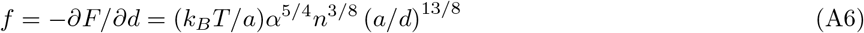

where *f*_*_ = *k*_*B*_*T/a* ≈ 1 pN, is a critical force which completely stretches the entropic degrees of freedom of the backbone chromatin. The correlation length induced by the stretching tension: *ξ*_*f*_ = *k*_*B*_*T/f*, is at a minimum under the critical force: *ξ*_*f*_(*f*_*_) ≈ *a*.

#### f. Monomer density along the backbone

The volume fraction of the monomers along the backbone *ϕ*_0_, is given by,

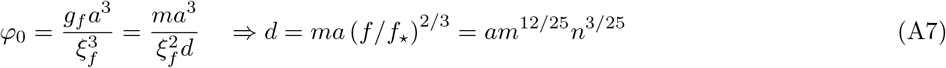

where *g*_*f*_ is the number of monomers in a force-induced blob: 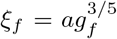. The number of monomers per backbone segment between two loops is *m* and the linear distance between two adjacent loop anchors is *d*, which we call the interloop distance.. As a simplifying step we consider monovalent loops while computing the steady-state interloop distance from the above force balance between loop repulsion and backbone stretching. Note that *d* is the extension of the polymer under tension *f*, which is also called the Pincus regime of polymer extension [80].

To keep our calculations simple, we will employ the limit *n* » *m*, which essentially implies that the chromosomes are always in the “polymer brush” regime where adjacent loops overlap with a varying degree. This contrasts the state where the backbone polymer is long and does not enforce loop overlap (*d* > *am*^3/5^), which is similar to the “random coil” or “unextruded” state and corresponds to an ordinary semidilute solution.

#### g. Dense axial core

The monomer volume fraction is maximum: *ϕ*_0_ ≈ 1, when the correlation length is equal to the monomer size: *ξ*_*f*_ = *a*, which can be established under a critical stretching tension *f*_*_ = *k*_*B*_*T/a* ≈ 0.4 pN. This leads to a densely packed core along the cylindrical backbone. However, the core at this stage is minimally thick *r*_0_ ≈ *a*, where *r*_0_ is the radius of the core. The thickness of the core may be increased by increasing the loop valency *α*.

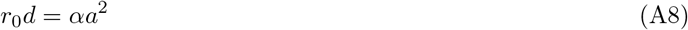

The above relationship between the core radius and loop valency is obtained from the condition that the lateral surface of the right-circular cylindrical core, ≈*r*_0_*d*, is saturated by the radially emanating *α* subloops.

Maximum core thickness corresponds to the case where the entire chromosome cross-section forms a compact core, which occurs for a loop valency *α*_*max*_. Using the condition: 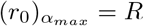, we get:

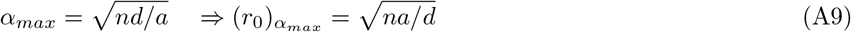

#### h. Persistence length of brush chromosomes

A thermally-excited bend generates a curvature *κ* along the cylindrical brush axes, that has a convex and a concave side. The volume accessible to the loops in the concave (convex) side is smaller (larger) than the unperturbed case by a factor of *κR* » 1. This leads to a perturbed volume fraction: 〈*ϕ*〉 (1 ± *κR*), where the upper/lower signs are for the concave/convex sides respectively, and 〈*ϕ*〉 = *na*^3^/(*R*^2^*d*) is the average unperturbed volume fraction inside the brush.

The free energy of a loop depends on the average volume fraction as, *F* = *k*_*B*_*Tn* 〈*ϕ*〉^5/4^ [(A5)]. The perturbation energy due to a curvature *κ* for a cylindrical brush with persistence length *ρ* is given by, *k*_*B*_*Tρκ*^2^*d*. Hence, we get the persistence length [39, 41]:

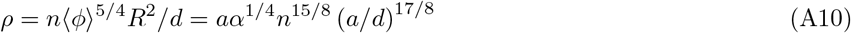

The above expression of the brush persistence length can also be consistently derived from the general relation between elastic moduli and persistence length using the following formula [41]: *ρ* = *R*^2^*d*(*∂*^2^*F/∂d*^2^).

##### Contribution from the core

The core behaves as a solid with an elastic modulus ≈ *k*_*B*_*T/a*^3^, and the corresponding persistence length depends on core thickness: 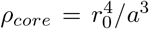. The core makes the chromosome stiffer, and the net persistence length of chromosomes may be obtained by adding the above contribution from the core to that of the loops. However, in the limit of saturated chromosomes, both the contributions have an identical scaling: *ρ* = *a*(*na/d*)^2^ [34]. Hence, the contribution from overlapping loops alone sufficiently accounts for chromosome stiffness, and we will employ (**??**) for persistence length in our calculations.

#### i. Fully stretched backbone for the optimal loop configuration

Overlap between optimal loops stretch the backbone where the transition to a fully stretched backbone occurs at the critical force value *f*_*_. For optimal loops, this critical repulsive force is obtained when 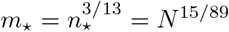.

### 2. Structure and mechanical rigidity of optimal chromosomes

#### a. Contour length

We obtain the axial contour length of cylindrical brush chromosomes with optimal loops and a fully stretched backbone, as follows:

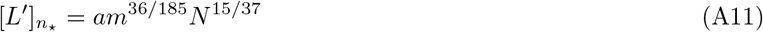

#### b. Thickness or radial extension

The average thickness of a chromatid is given by,

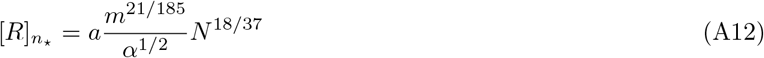

#### c. Core width

The diameter of the core scales positively with the level of saturation in the following way.

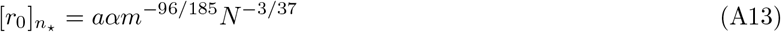

#### d. Persistence length

The brush persistence length obtained from (A10) for an optimal configuration is given as follows:

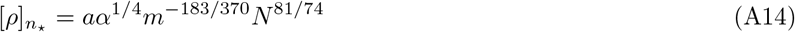

#### e. Doubling force

The extrapolated Hookean force associated with doubling the length of chromosomes is an intensive quantity that can be measured experimentally [19, 47], and which we define within our model as: *f*_0_ = *d*(*∂*^2^*F/∂d*^2^). For an optimal configuration, we have the following for the doubling force.

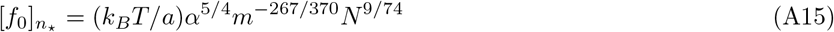

#### f. Entropic repulsion between sister chromatids

The origin of entropic repulsion between the sister chromatid arms is the high osmotic pressure developed in the volume where the sister chromatids overlap. The repulsive force may be computed from the net osmotic pressure over the cross-sectional area of overlap between the intertwined sister chromosomes, ≈ *RL′*.

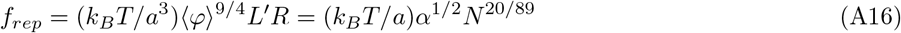

Note that the repulsive force is ≈ 10 pN for monovalent optimal loops of human chromosomes, which drives physical segregation of the intertwined sister chromosomes, a result of the polymer brush morphology of the sister chromatids.

#### g. Elastic modulus

The elastic moduli of chromosomes may be obtained from the average volume fraction of nucleosome monomers inside the cylindrical brush in the following manner [1].

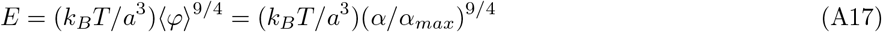

where *k*_*B*_*T/a*^3^ ≈ 4 kPa, is the maximum elastic modulus of chromosomes in the limit of the core spanning the entire chromosome.

Metaphase chromosomes of higher eukaryotes are known to have an elastic modulus *E*_*meta*_ ≈ 1 kPa [45]. The value of valency required to generate *E*_*meta*_ scales with the total chromosome length, and is given by *α*_*_ = (0.5)*N* ^40/89^. Note that *α*_*_ corresponds to a completely stretched backbone configuration (*m* = *m*_*_).

##### a. Prophase limit (fully stretched backbone)

In case of a fully stretched backbone, i.e., *m* = *m*_*_, the brush becomes stiffer and the above scalings slightly change:

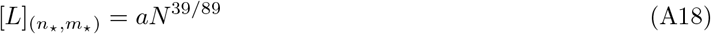

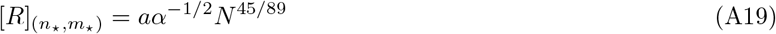

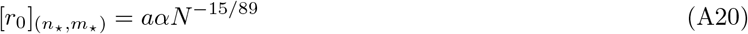

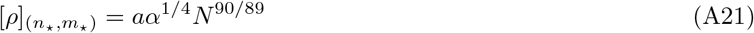

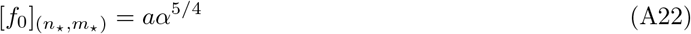

##### b. Metaphase limit (highly branched loops)

In metaphase, the valency *α* = *α*_*_ = *N* ^40/89^ makes the brush even more stiff and compact and further modifies the scaling:

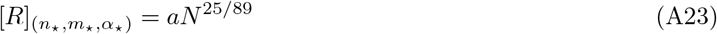

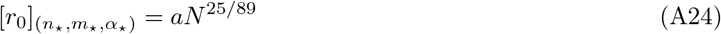

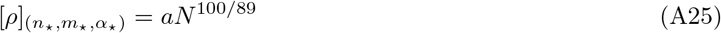

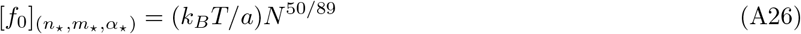

### 3. Confined solution of chromosomes

We model the nuclear confinement of chromosomes by defining an average volume fraction of the confined genome *φ*, which introduces a correlation length associated with the confinement volume fraction.

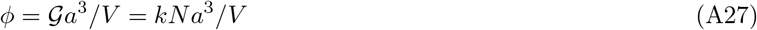

where 𝒢 and *N* are, respectively, the total number of monomers in the genome and the chromosome; and *k* is the total number of chromosomes or the karyotype of the cell: 𝒢 = *kN*. The volume of the nucleus is denoted by *V* [Table II]. Note that chromosomes inside nuclear confinement are expected to have a high degree of overlap: *kR_F_* (*N*)^3^ » *V*.

**TABLE II.** Genome size in a diploid nuclei in Mega-base pair units (1 Mb=10^3^ kb) and karyotype *k* are used to obtain the average chromosome length *N*. The number of monomers or nucleosomes may be obtained by dividing the chromosome length in Mb by *≈* 0.2 kb corresponding to one nucleosome. Nuclear volumes of various organisms *V* are used to compute the average volume fraction of chromatin inside nuclear confinement: *φ* = *kNa*^3^/*V*, where *a* ≈ 10 nm is the nucleosome diameter. Note, bacterial chromosomes are made up of cylindrical segments of length *a* ≈_2_ 50 nm and width *b* ≈ 5 nm (corresponding to protein-bound DNA), where the volume fraction is computed as *φ* = *kNab*^2^/*V*.

| organism | genome size<br>(Mb) | karyotype<br>$k$ | chromosome<br>size (Mb) | $N$ | nuclear volume<br>$V$ ($\mu\text{m}^3$) | volume fraction<br>$\phi$ |
| --- | --- | --- | --- | --- | --- | --- |
| Lily (*L. longiflorum*) | 97000 | 24 | 4000 | $2 \times 10^7$ | 1347 [82] | 0.36 |
| Newt (*N. viridescens*) | 95000 | 24 | 4000 | $2 \times 10^7$ | 4174 [83] | 0.11 |
| Human (*H. sapiens*) | 6000 | 46 | 130 | $6 \times 10^5$ | 300 [83, 84] | 0.1 |
| Mouse (*M. musculus*) | 5600 | 40 | 140 | $7 \times 10^5$ | 400 [83, 85, 86] | 0.07 |
| Toad (*X. laevis*) | 5400 | 46 | 150 | $5.9 \times 10^5$ | 307 [83] | 0.09 |
| Chicken (*G. gallus*) | 2200 | 78 | 28 | $1.4 \times 10^5$ | 210 [83] | 0.05 |
| Fly (*D. melanogaster*) | 280 | 8 | 35 | $1.7 \times 10^5$ | 78 [83] | 0.02 |
| Thale cress (*A. thaliana*) | 240 | 10 | 24 | $1.2 \times 10^5$ | 70 [87, 88] | 0.02 |
| Nematode (*C. elegans*) | 200 | 12 | 17 | $8.3 \times 10^4$ | 200 [89, 90] | 0.005 |
| Fission yeast (*S. pombe*) | 25 | 6 | 4 | $2.1 \times 10^4$ | 12 [91–93] | 0.01 |
| Budding yeast (*S. cerevisiae*) | 25 | 32 | 0.8 | $3.9 \times 10^3$ | 3 [83, 94, 95] | 0.04 |
| Bacteria (*E. coli*) | 9.2 | 2 | 4.6 | $3 \times 10^4$ | 2 [96] | 0.02 |

**TABLE III.** Chromosome loop sizes for various organisms corresponding to interphase (TAD) and mitosis are tabulated. Experimental values for the structure and mechanical properties: chromosome axial contour length *L′*, width *R*, doubling force *f*_0_ in nano-Newton units (nN), and elastic modulus or Young’s modulus *E* in kilo-Pascal units (kPa). Note, the elastic modulus reported in Ref. [98] is for migratory grasshopper (*M. sanguinipes*) chromosomes, however, due to lack of genomic data on *M. sanguinipes*, we use the genomic data for meadow grasshopper (*C. parallelus*).

| organism | chromosome size (Mb) | chromosome domains and TADs (kb) | metaphase loops (kb) | $L'$ ($\mu\text{m}$) | $R$ ($\mu\text{m}$) | $f_0$ (nN) | $E$ (kPa) |
| --- | --- | --- | --- | --- | --- | --- | --- |
| Newt (*N. viridescens*) | 4000 | — | — | 20 [48] | 1.6 [48] | 1 [48] | 0.7 [48] |
| Pine (*P. taeda*) | 1800 | — | — | 18 [97] | 1 [97] | — | — |
| Grasshopper (*C. parallelus*) | 1400 | — | — | — | — | — | 0.43 [98] |
| Deer (*M. muntjak*) | 750 | — | — | 11 [99] | 0.96 [99] | — | — |
| Barley (*H. vulgare*) | 729 | — | — | 12 [97] | 0.9 [97] | — | — |
| Toad (*X. laevis*) | 150 | — | 90 [100] | 5 [48, 100] | 0.8 [48] | 0.6 [47] | 0.4 [48] |
| Mouse (*M. musculus*) | 140 | 350 [29], 880 [101] | 50 [102] | — | — | — | — |
| Human (*H. sapiens*) | 130 | 200 [32], 880 [101] | 50 [9] | 4.3 [99] | 0.88 [99] | 0.25 [19] | 0.42 [19] |
| Zebrafish (*D. rerio*) | 78 | 500 [103] | — | — | — | — | — |
| Fly (*D. melanogaster*) | 35 | 60 [101, 104, 105] | — | 3.9 [97] | 0.4 [97] | — | — |
| Rice (*O. sativa*) | 31 | 45 [106], 487 [107] | — | 2.5 [97] | 0.3 [97] | — | — |
| Chicken (*G. gallus*) | 28 | 80 [25] | — | — | — | — | — |
| Roundworm (*C. elegans*) | 17 | 1000 [108] | — | — | — | — | — |
| Fission yeast (*S. pombe*) | 4 | 50 [109, 110], 80 [111, 112] | — | — | — | — | — |
| Budding yeast (*S. cerevisiae*) | 0.8 | 100 [113], 200 [114] | — | 1 [115] | 0.5 [115] | — | — |
| Bacteria (*E. coli*) | 4.6 | 170 [116] | — | 2 [55] | — | — | — |

#### a. Entanglements in confinement

We estimate inter-chromosome entanglement from the number of nearby contacts between different chromosomes. Since the semidilute solution of chromosomes may be viewed as a closely packed system of blobs, the total number of inter-blob collisions, which scales with the total number of blobs, gives the level of entanglement in the system. However, note that we are treating the chromosome as a renormalized polymer with segment size *R* and a persistence length *ρ* > *R*, such that the number of segments per chromosome ≈ *L′*/*R* ≡ *N′* » *N*.

Each of these cylindrical brush segments have a volume *v* ≈ *R*^2^*ρ*, and an excluded volume *w* ≈ *ρ*^2^*R*. The correlation length in confinement is given by, 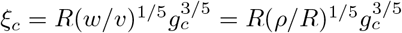 [1].

The number density of chromosome monomers in the confined volume is uniform:

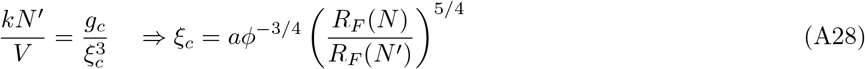

where *R*_*F*_ (*N′*) = *R*(*w/v*)^1/5^(*N′*)^3/5^, is an equilibrium length scale denoting the end-to-end distance of the chromosome polymer in the non-overlapping or dilute limit. Note that the above equation is applicable to the semidilute regime, it is only expressed in a dilute-regime length scale for notational convenience. The total number of confinement blobs, given as follows,

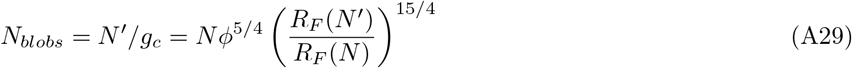

that scales positively with the chromatin volume fraction *φ* [6]. This indicates a higher entanglement between chromosomes when the average nucleosome concentration in the nucleus is higher.

The level of inter-chromosome entanglement, which we denote by average catenation-squared 〈Ca^2^〉, since every catenation irrespective of their sign contribute to entanglement, a consequence of fluctuating topology, scales with the number of blobs, we get the following for inter-chromosomal entanglements per chromosome.

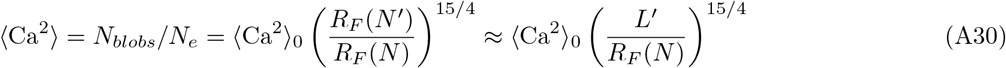

where

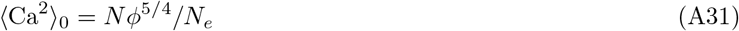

is the level of entanglement in a semidilute chromatin solution, i.e., in the unextruded state; and *N*_*e*_ ≈ 100 is the entanglement number which is a constant of proportionality [6, 43, 81].

For the optimal configuration, the renormalized chromosome polymer behaves as a semiflexible object: *N′* = *L′*/*R* ~ 1, in which case we may write: *R*_*F*_(*N′*) ≈ *L′*, indicating that optimal loop size minimizes both chromosome axial length and inter-chromosomal entanglements.

Inter-chromosome entanglements per chromosome, for an optimal configuration is given by,

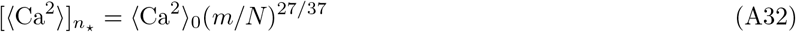

#### b. Ideal or Gaussian polymer behavior of semiflexible chromosomes

Optimal chromosomes exhibit semiflexible behavior, i.e., *ρ* > *R*. It has been argued that semiflexible polymers in semidilute regimes may show theta-solvent behavior, i.e. Gaussian statistics or ideal polymer statistics [44]. In such a case, the end-to-end distance of an optimal chromosome polymer is expected to be: *R*_Θ_(*N′*) = (*RρN′*)^1/2^.

Two neighboring (e.g., tethered [6]) Gaussian phantom chains of *N* segments each have inter-chain linking-squared scale as ~ *N* ^1/2^. For Gaussian-polymer in confinement, the net linking or catenation-squared per chromosome is proportional to the volume fraction of each chromosome polymer in confinement, which denotes the probability of contact, times the catenation for two neighbor chains [7]:

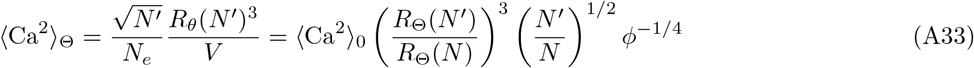

where, 〈Ca^2^〉_0_ is the entanglement in semidilute (self-avoiding) linear chromatin, given by (A31); and 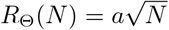 is an equilibrium length scale for chromatin chain. Fig. 5 shows 〈Ca^2^〉_Θ_ as a contour shade for various biologically relevant *N* and *φ*, where we have used *R*_Θ_(*N′*) ≈ *L′*, because *N′* ~ 1 for optimal chromosomes. Importantly, Fig. 5 shows that the scalings in (A33) and (A30) are not much different (compare with Fig. 3 in the main text). As a result, our conclusion: compaction of brush morphology drives chromosome disentanglement, is immune to whether we use semiflexible or flexible polymer scaling for chromosomes. This is derived from the fact that the renormalized chains are short (about one statistical segment long) where the semiflexible nature does not play a noticeable role.

### 4. Bacterial chromosomes: cylindrical DNA monomers

Bacteria does not have nucleosomes, however, bacterial DNA is covered with various DNA binding proteins that slightly increases the thickness of the DNA cross-section from its bare cross section of ≈ 2 nm. We consider bacterial DNA to be a polymer of cylindrical monomers of height *a* = 50 nm, which is the persistence length of bare DNA, and diameter *b* = 5 nm corresponding to protein-bound DNA.

**FIG. 5.**
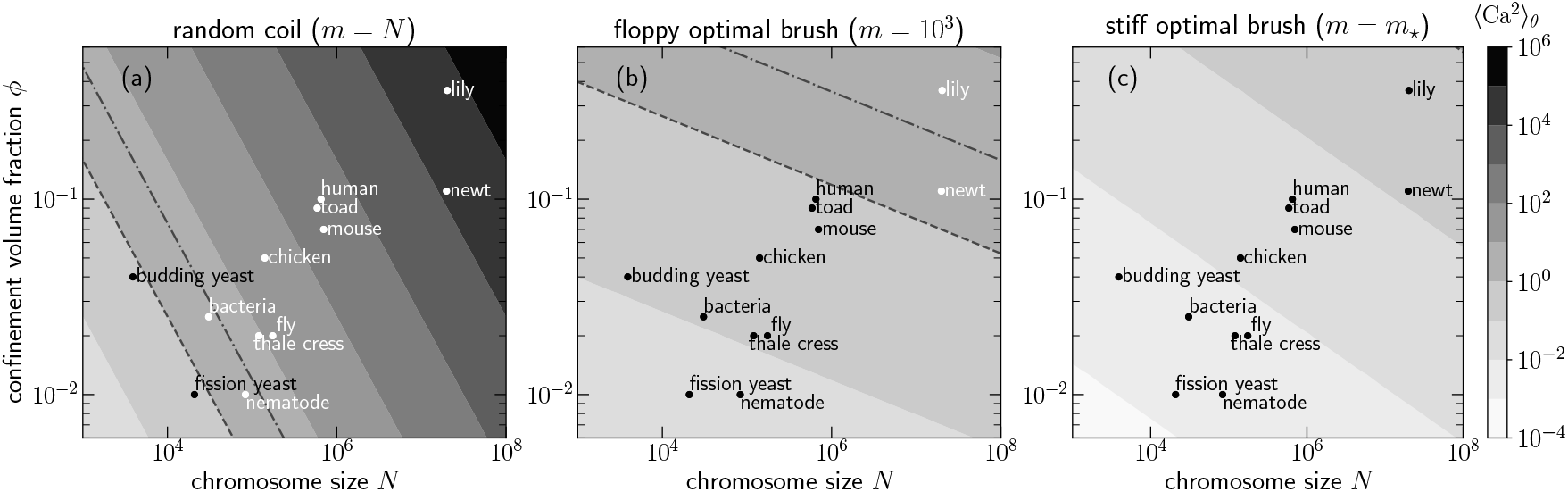
Inter-chromosome entanglement per chromosome, treating the chromosomes as phantom semiflexible polymers that exhibit Gaussian polymer statistics (A33). Fig. (a) shows the entanglement in unfolded chromatin in the semidilute limit (A31). Compare (b) and (c) with respectively Figs. 3(b) and 3(c) of the main text where renormalized chromosomes were considered to be flexible polymers (A30). This shows that irrespective of whether we posit, brush chromosomes, due to their semiflexible nature show weak self-avoidance and behave like Gaussian polymers [44], or they are flexible self-avoiding polymers, compaction-driven segregation and global topological disentanglement is expected for the biologically relevant range of polymer sizes and confinement.

The monomer aspect ratio, defined as the ratio of monomer excluded volume to monomer volume: *w/v* = (*a*^2^*b*)/(*ab*^2^) ≈ 10 for bacterial chromatin. The equilibrium end-to-end distance of unconfined bacterial genome: *R*_*F*_ (*N*) = *b*(*w/v*)^1/5^*N* ^3/5^ Following the steps outlined above, i.e., from Eq. A2 to Eq. A7, we find *R* (*w/v*)^1/4^ and *d* ~ (*w/v*)^1/12^. This when plugged into the minimization of the axial contour length generates a very weak dependence for the optimal loops: *n*_*_ ~ (*w/v*)^0.08^. The contour length for optimal chromosomes scales as *L′*(*w/v*)^0.25^, which is within a factor-of-two for the contour length of bacterial chromosomes and does not introduce significant scaling. The parameter that affects entanglement has a very weak scaling: *L′*/*R*_*F*_ (*N*) ∼ (*w/v*)^0.05^. As a result, our discussion on flexible eukaryotic chromatin (where (*w/v*) *≈* 1) is also applicable to the bacterial case.

